# Regeneration and Developmental Enhancers Are Differentially Compatible with Minimal Promoters

**DOI:** 10.1101/2022.06.20.496839

**Authors:** Ian J. Begeman, Benjamin Emery, Andrew Kurth, Junsu Kang

**Affiliations:** Department of Cell and Regenerative Biology, School of Medicine and Public Health, University of Wisconsin–Madison, Madison, WI, 53705, USA.; UW Carbone Cancer Center, School of Medicine and Public Health, University of Wisconsin– Madison, Madison, WI, 53705, USA

## Abstract

Enhancers and promoters are *cis*-regulatory elements that control gene expression. Enhancers are activated in a cell type-, tissue-, and condition-specific manner to stimulate promoter function and transcription. Zebrafish have emerged as a powerful animal model for examining the activities of enhancers derived from various species through transgenic enhancer assays, in which an enhancer is coupled with a minimal promoter. However, the efficiency of minimal promoters and their compatibility with multiple developmental and regeneration enhancers have not been systematically tested in zebrafish. Thus, we assessed the efficiency of six minimal promoters and comprehensively interrogated the compatibility of the promoters with developmental and regeneration enhancers. We found that the *fos* minimal promoter and *Drosophila* synthetic core promoter (DSCP) yielded high rates of leaky expression that may complicate the interpretation of enhancer assays. Notably, the adenovirus *E1b* promoter, the zebrafish *lepb* 0.8-kb (*P0.8*) and *lepb* 2-kb (*P2*) promoters, and a new zebrafish synthetic promoter (*ZSP*) that combines elements of the *E1b* and *P0.8* promoters drove little or no ectopic expression, making them suitable for transgenic assays. We also found significant differences in compatibility among specific combinations of promoters and enhancers, indicating the importance of promoters as key regulatory elements determining the specificity of gene expression. Our study provides guidelines for transgenic enhancer assays in zebrafish to aid in the discovery of functional enhancers regulating development and regeneration.

## INTRODUCTION

Enhancers are noncoding *cis*-regulatory elements that serve as vital regulators of spatiotemporal gene expression (Panigrahi and O’Malley, 2021; Shlyueva et al., 2014; Tippens et al., 2018). Enhancers selectively activate gene expression in a highly regulated manner in various conditions and developmental stages (Jindal and Farley, 2021; Rickels and Shilatifard, 2018). Thus, identifying and characterizing enhancers in particular tissues and contexts will expand our understanding of the regulatory mechanisms governing essential biological phenomena, such as development and regeneration.

Enhancer candidates can be identified through genome-wide assays of accessible chromatin regions via assay for transposase-accessible chromatin using sequencing (ATAC-seq) or through chromatin immunoprecipitation followed by sequencing (ChIP-seq) for assaying specific enhancer-associated histone modifications, such as histone H3 lysine 27 acetylation (H3K27ac) (Barski et al., 2007; Bonn et al., 2012; Buenrostro et al., 2013; Shlyueva et al., 2014). While the presence of open chromatin and specific histone marks is suggestive of active enhancers, the in vivo activity of enhancer candidates must be verified empirically (Cao et al., 2022; Goldman et al., 2017). A widely used in vivo method is the transgenic reporter assay, in which an enhancer is paired with a minimal promoter and a reporter gene and then inserted into the genome of a model organism, with reporter gene expression providing a readout of the enhancer’s activity (Begeman et al., 2020b; Dickel et al., 2016; Dobrzycki et al., 2020; Goldman et al., 2017; Hewitt et al., 2017; Kang et al., 2016; Kvon, 2015; Osterwalder et al., 2018; Wong et al., 2020). Because promoters are also important regulators of gene expression, a minimal promoter with little or no background expression is required to obtain conclusive results from transgenic enhancer assays.

Zebrafish (*Danio rerio*) have emerged as a powerful system for examining enhancers, owing to their genetic tractability, transparency in early developmental stages, and high fecundity (Kang et al., 2016; Yuan et al., 2018). Enhancers derived from other species, such as humans, mice, and sponges, can also direct conserved expression patterns in zebrafish, highlighting the usefulness of zebrafish for cross-species enhancer investigation (Heller et al., 2022; Perez-Rico et al., 2017; Wong et al., 2020; Yuan et al., 2018). Recent genome-wide analyses of zebrafish have rapidly expanded the list of enhancer candidates that may contribute to development and regeneration (Cao et al., 2022; Goldman et al., 2017; Kang et al., 2016; Quillien et al., 2017). However, published enhancer studies in zebrafish have employed a variety of minimal promoters (**Table S1**), and no promoter has yet been accepted as a common standard by the zebrafish community. Moreover, the leakiness of these minimal promoters and their compatibility with distinct classes of enhancers have not been systematically determined.

In this study, we assessed the efficiency of six minimal promoters and comprehensively interrogated their compatibility with developmental and regeneration enhancers. We found that two minimal promoters yielded high rates of leaky expression that may complicate the interpretation of enhancer assays. Notably, four minimal promoters drove little or no ectopic expression, making them suitable for transgenic assays. We also demonstrated significant differences in compatibility among specific combinations of promoters and enhancers. Our study provides guidelines for transgenic enhancer assays in zebrafish to aid in the discovery of functional enhancers regulating development and regeneration.

## RESULTS

### Minimal promoters drive different rates of the leaky expression

We performed a literature search of enhancer assays and selected five minimal promoters to test for efficiency in transgenic enhancer assays in zebrafish: *E1b*, *leptin b (lepb)* 0.8 kb (*P0.8)*, *lepb* 2 kb (*P2)*, *fos*, and the *Drosophila* synthetic core promoter (DSCP). The *E1b* promoter, referred to as *E1b* hereafter, originates from the adenovirus *early region 1b* gene and has been widely used for validating developmental enhancers in zebrafish (**Table S1**) (Argenton et al., 1996; Bar Yaacov et al., 2019; Birnbaum et al., 2012; Booker et al., 2013; Hirsch et al., 2018; Kim et al., 2014; Laarman et al., 2019; Li et al., 2010; Oksenberg et al., 2014; White, 2001; Yanovsky-Dagan et al., 2015). *P0.8* and *P2* are the 793-bp and 2045-bp sequences, respectively, immediately upstream of the *lepb* transcription start site (Kang et al., 2016). *lepb* shows no detectable developmental expression in the heart or fins but is strongly upregulated in those tissues during regeneration. A *lepb-*linked regeneration enhancer known as *LEN* is located 5.2 kb upstream of *lepb* and is required for regeneration-dependent *lepb* induction (Kang et al., 2016; Thompson et al., 2020). *P0.8* and *P2* have been used as basal promoters for identifying and testing regeneration enhancers, including *cardiac LEN (cLEN*), which is the enhancer fragment of *LEN* that directs expression in injured hearts (Begeman et al., 2020b; Geng et al., 2021; Kang et al., 2016; Karra et al., 2018; Lee et al., 2020). The 100-bp *fos* promoter, referred to as *fos* hereafter, is derived from the mouse *fos* gene and has been used in zebrafish for studying both developmental and regeneration enhancers (Cao et al., 2022; Goldman et al., 2017; Scott and Baier, 2009; Thompson et al., 2020). The 155-bp DSCP is an optimized synthetic core promoter that has been utilized for examining neuronal enhancers and for performing large-scale developmental enhancer screens in *Drosophila* (Kvon et al., 2014; Pfeiffer et al., 2008).

An important characteristic of a minimal promoter is having limited intrinsic leaky activity, as such activity can hinder the interpretation of the activity of the enhancer being tested. To determine the leakiness of the selected minimal promoters, we generated constructs containing *E1b*, *P0.8*, *P2*, *fos*, and DSCP upstream of EGFP in the absence of an enhancer. We assessed the basal activity of these constructs by microinjecting them into one-cell-stage zebrafish embryos and examining EGFP expression during larval development (**Fig. 1A**). The constructs contained a lens-specific *mCherry* reporter gene that enables the selection of transgenic larvae. We found that *E1b*, *P0.8*, and *P2* yielded low rates of background expression with no detectable EGFP expression in the heart, eye, or fin fold of larvae (n=196, 57, and 88, respectively) (**Fig. 1B, C and Table S2**). In contrast, *fos* and DSCP yielded high rates of leaky expression (**Fig. 1B, C and Table S2**). *fos* drove expression in the heart, eye, and tail fin fold in 10%, 2%, and 78% of larvae, respectively (n=134). Among larvae carrying the DSCP construct, 43%, 3%, and 100% directed EGFP expression in the heart, eye, and tail fin fold, respectively (n=63). These results suggest that *E1b*, *P0.8*, and *P2* may be preferable to *fos* and DSCP due to their lower basal activity.

**Figure 1.**
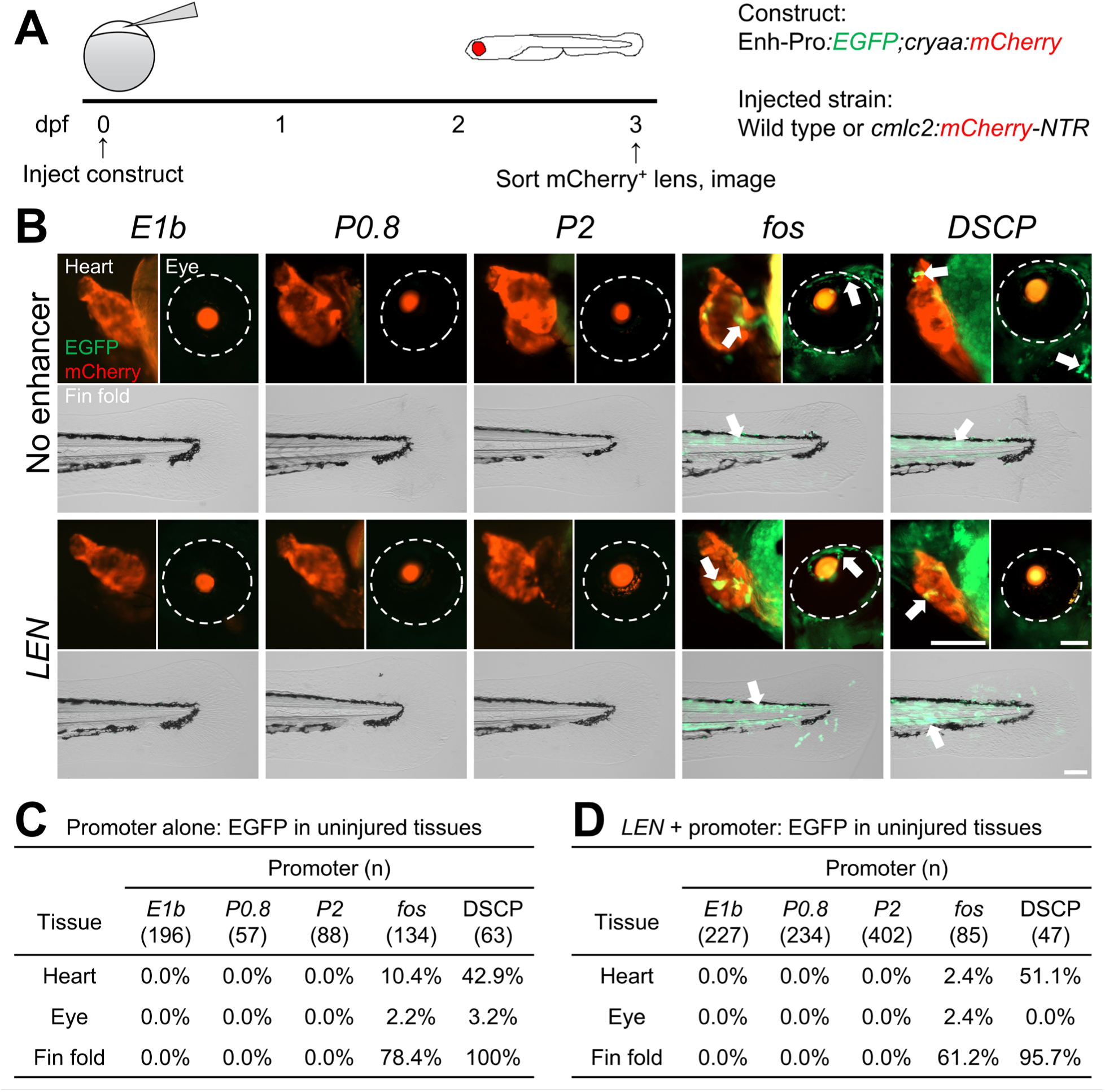
*E1b*, *P0.8*, and *P2* drive minimal leaky expression in uninjured tissues. (**A**) Experimental design. Transgenic constructs were injected into one-cell-stage wild-type or Tg(*cmlc2:mCherry-NTR*) embryos, and EGFP expression was observed at 3 dpf. (**B**) 3-dpf F0 uninjured hearts, eyes, and fin folds. Arrows indicate EGFP expression. Eyes are demarcated by dotted lines. (**C, D**) Tables of larvae displaying EGFP expression in each tissue for each promoter alone (**C**) or when paired with *LEN* (**D**). Scale bars: 100 μm.

We next determined whether placing an enhancer fragment upstream of the minimal promoters influences their activity in larvae (**Fig. 1A**). We selected *LEN*, a well-characterized regeneration enhancer that exhibits no activity during development or in uninjured tissues but drives injury-responsive expression in the heart and fin (Kang et al., 2016). Because *LEN* activity is regeneration-specific, we expected to observe no detectable expression in uninjured larvae if the promoter exhibited no leaky expression. We observed limited basal activities of *LEN-E1b*, *LEN-P0.8*, and *LEN-P2*, as no larvae expressed EGFP in the heart, eye, or fin fold (n=227, 234, and 402, respectively) (**Fig. 1B, D and Table S2**). In contrast, a substantial number of *LEN-fos* and *LEN-DSCP* larvae displayed leaky expression throughout the body (**Fig. 1B, D and Table S2**). A total of 2%, 2%, and 61% of *LEN-fos* larvae expressed EGFP in the heart, eye, and tail fin fold, respectively (n=85), while 51%, 0%, and 96% of *LEN-DSCP* larvae drove expression in the heart, eye, and tail fin fold, respectively (n=47). The low levels of background expression of *E1b, P0.8*, and *P2* indicate that they can serve as adequate minimal promoters for F_0_ transgenic enhancer assays in zebrafish. In contrast, the high rates of leaky activity of *fos* and DSCP may mask the activity of paired enhancers and thus complicate conclusions drawn from enhancer assays using F_0_ mosaic embryos.

### Developmental enhancers display preferences for specific promoters

Another important feature of minimal promoters is the ability to drive expression at the locations and timepoints specified by the paired enhancer. We sought to determine how efficiently *E1b*, *P0.8*, and *P2* direct expression in association with developmental enhancers. We placed each promoter downstream of two validated developmental enhancers: *103runx1EN* and *gata2a-i4* (**Fig. 2A**)*. 103runx1EN* is located 103 kb upstream of the zebrafish *runx1* gene and displays multiple activities depending on the tissues involved and injury status (Goldman et al., 2017). *103runx1EN* directs expression in the notochord and CNS of uninjured developing larvae and in the heart and fin in response to injury, indicating that it can function as either a developmental or regeneration enhancer. *gata2a-i4* is an evolutionarily conserved enhancer that regulates the developmental expression of *gata2a* in endothelial cells (Dobrzycki et al., 2020; Gao et al., 2013). In agreement with previous reports (Dobrzycki et al., 2020; Gao et al., 2013; Goldman et al., 2017), our F_0_ transgenic enhancer assays revealed that *103runx1EN* drove EGFP expression in the notochord and CNS but not the heart, eye, or fin fold (**Fig. 2B, C and Table S2**). However, the expression rate of each *103runx1EN*-promoter construct varied significantly, with notochord expression being detectable in 15% of *103runx1EN-E1b*, 1% of *103runx1EN-P0.8*, and 6% of *103runx1EN-P2* larvae (n=414, 246, and 166, respectively). The expression rates driven by *gata2a-i4* also varied significantly among promoters (**Fig. 2B, D and Table S2**). Expression was observed in the heart and vasculature of 77% and 95% of *gata2a-i4-E1b* larvae, respectively (n=168). In contrast, only 2% of *gata2a-i4-P0.8* larvae expressed EGFP in the heart or vasculature (n=61). No detectable cardiac or vasculature expression was observed in *gata2a-i4-P2* larvae (n=73). These data suggest that minimal promoters exhibit different degrees of compatibility with developmental enhancers in F_0_ transgenic enhancer assays. Among the tested minimal promoters, our results indicate that *E1b* exhibits the most efficient compatibility with developmental enhancers.

**Figure 2.**
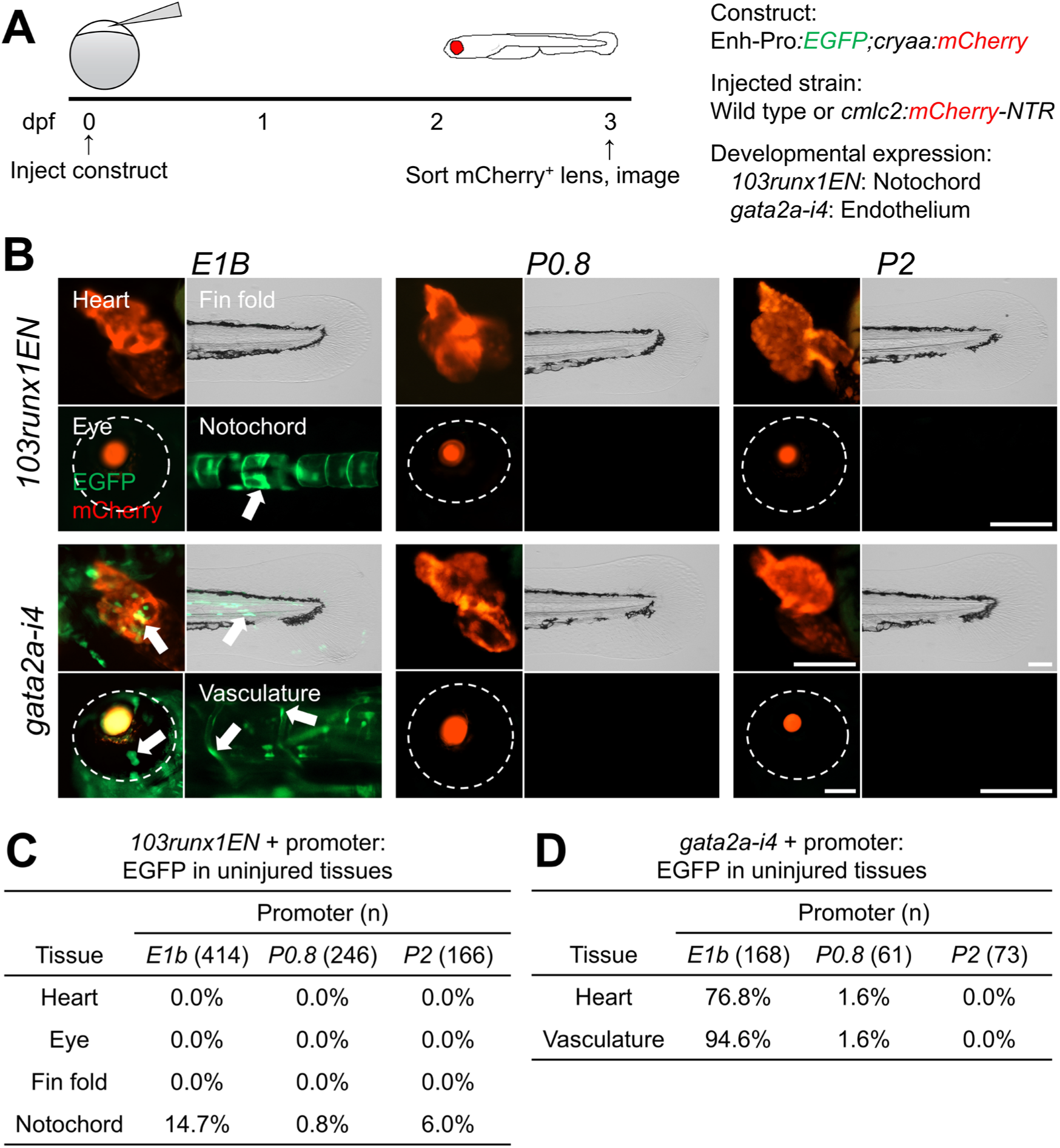
Developmental enhancer activity is preferentially compatible with *E1b*. (**A**) Experimental design. Transgenic constructs were injected into one-cell-stage wild-type or Tg(*cmlc2:mCherry-NTR*) embryos, and EGFP expression was observed at 3 dpf. (**B**) 3-dpf F0 hearts, eyes, fin folds, notochords, and vasculature in uninjured larvae. Arrows indicate EGFP expression. Eyes are demarcated by dotted lines. (**C, D**) Tables of larvae displaying EGFP expression in each tissue for each promoter when paired with *103runx1EN* (**C**) or *gata2a-i4* (**D**). Scale bars: 100 μm.

### Differential injury-induced expression among enhancer-promoter pairs

Regeneration enhancers are characterized as being quiescent in uninjured tissues but robustly activated upon injury (Begeman and Kang, 2018; Kang et al., 2016). Important features of the minimal promoters used for investigating regeneration enhancers include a lack of intrinsic injury-responsive activity and the ability to mediate injury-inducible activation of paired enhancers. To determine how tightly *E1b, P0.8*, and *P2* are regulated in injured contexts, we examined the activity of each promoter both alone and when paired with the regeneration enhancers *LEN* and *103runx1EN*. Previous studies validated the activities of these two regeneration enhancers in injured hearts and fins (Goldman et al., 2017; Kang et al., 2016). To selectively injure the hearts of larvae, we employed the *cardiac myosin light chain 2* (*cmlc2*)*:mCherry-nitroreductase* (*NTR)* system, which enables the genetic ablation of cardiomyocytes (CMs) (**Fig. 3A**). Under the control of the *cmlc2* promoter, the bacterial *NTR* gene is expressed specifically in CMs. When larvae are treated with the innocuous prodrug metronidazole (Mtz), NTR converts Mtz into a cytotoxic molecule, resulting in the ablation of a substantial number of CMs (Curado et al., 2007; Curado et al., 2008; Dickover et al., 2013). We observed little or no cardiac expression in *E1b*, *P0.8*, and *P2* larvae at 3 days post-treatment (dpt) (2%, n=154; 0%, n=96; and 0%, n=109, respectively), indicating minimal leaky induction of these promoters following injury (**Fig. 3B, C and Table S2**). When paired with cardiac regeneration enhancers, *E1b*, *P0.8*, and *P2* displayed different degrees of compatibility. *LEN* drove cardiac expression in 18%, 61%, and 42% of larvae when paired with *E1b*, *P0.8*, and *P2,* respectively (n=74, 61, and 139) (**Fig. 3B, C and Table S2**). *103runx1EN* displayed a similar but more biased pattern, driving cardiac EGFP expression in 5%, 48%, and 45% of larvae when paired with *E1b*, *P0.8*, and *P2*, respectively (n=270, 148, and 102) (**Fig. 3B, C and Table S2**). These data suggest preferential compatibility of cardiac regeneration enhancer activity with *P0.8* and *P2* over *E1b*.

**Figure 3.**
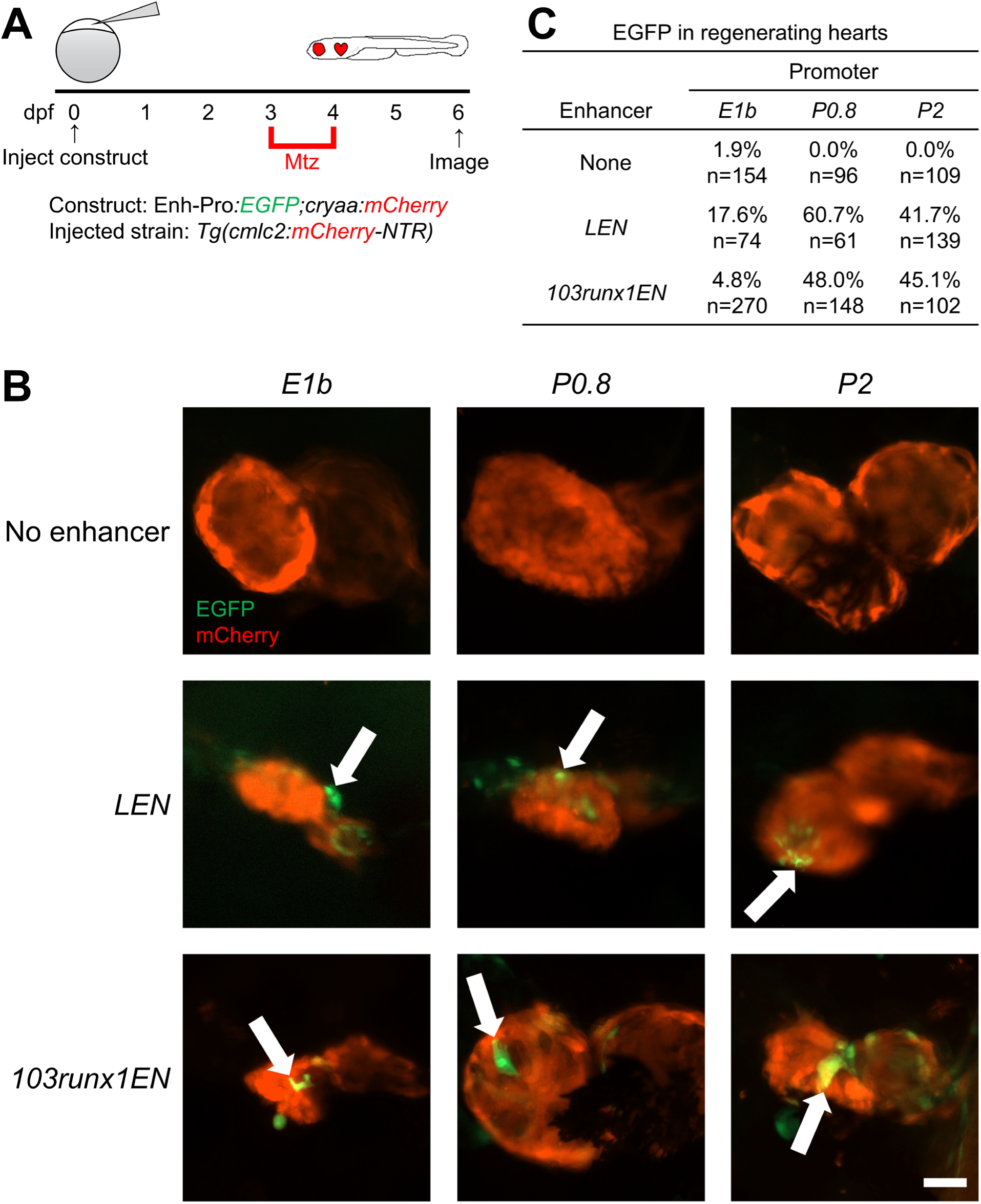
Regeneration enhancer activity in hearts is preferentially compatible with *P0.8* and *P2*. (**A**) Experimental design. Transgenic constructs were injected into one-cell-stage Tg(*cmlc2:mCherry-NTR*) embryos, larvae were treated with Mtz for 24 hours starting at 3 dpf to ablate cardiomyocytes, and EGFP expression was observed at 6 dpf. (**B**) 6-dpf F0 regenerating hearts following Mtz-induced cardiomyocyte ablation. Arrows indicate EGFP expression. (**C**) Table of larvae displaying cardiac EGFP expression for each promoter alone or when paired with *LEN* or *103runx1EN*. Scale bar: 50 μm.

We next utilized larval tail regeneration assays, which are a model for studying appendage regeneration (**Fig. 4A**). Amputating the distal tip of the tail removes multiple tissues, including the notochord, blood vessels, epithelia, and mesenchyme (Mateus et al., 2012; Yoshinari et al., 2009). Of note, we previously found that *P2,* but not *P0.8*, drove injury-induced expression in the larval tail, although this injury-inducible ability of *P2* alone was not observed in the amputated adult fins (Kang et al., 2016). Consistent with our previous reports, no *P0.8* larvae directed expression in the amputated larval tail (n=55), while 38% of *P2* larvae drove expression (n=56) (**Fig. 4B, C and Table S2**). Meanwhile, 4% of *E1b* larvae exhibited EGFP expression in the injured tail (n=124) (**Fig. 4B, C and Table S2**), suggesting that *E1b* possesses limited but evident injury-responsive activity. *E1b* and *P2* are predicted to contain injury-responsive elements, including transcription factor (TF) binding sites for AP-1, a TF that is important for regeneration (Beisaw et al., 2020) (**Fig. S1**). Given that *E1b* and *P2* do not direct injury-dependent expression in adult fins (**Fig. S2**), the injury-responsive elements are likely repressed during maturation.

**Figure 4.**
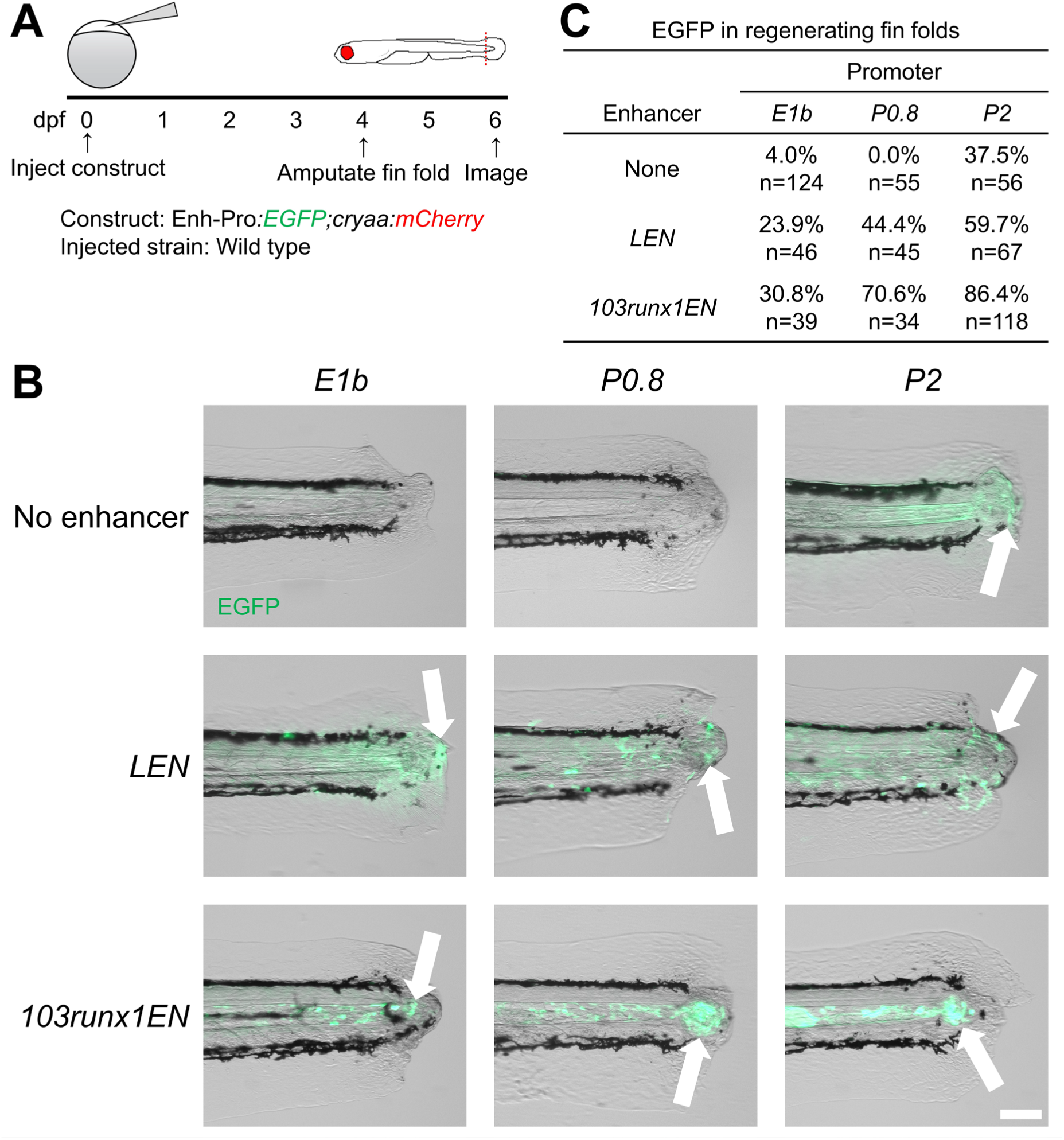
Injury-induced activity of enhancer-promoter constructs in amputated fin folds. (**A**) Experimental design. Transgenic constructs were injected into one-cell-stage wild-type embryos, fin folds were amputated at 4 dpf, and EGFP expression was observed at 6 dpf. (**B**) 6-dpf F0 regenerating fin folds following injury. Arrows indicate EGFP expression. (**C**) Table of larvae displaying injury-induced fin fold EGFP expression for each promoter alone or when paired with *LEN* or *103runx1EN*. Scale bar: 100 μm.

We next assessed minimal promoter activity in injured fins when paired with regeneration enhancers. For all tested minimal promoters, *LEN* and *103runx1EN* robustly increased the rate of EGFP expression in injured fins, although the efficiency differed between promoters. Injury-induced expression in the tail was observed in 24%, 44%, and 60% of *LEN-E1b*, *LEN-P0.8*, and *LEN-P2* larvae, respectively (n=46, 45, and 67) (**Fig. 4B, C and Table S2**). *103runx1EN* drove injury-induced expression in 31%, 71%, and 86% of larvae when paired with *E1b*, *P0.8*, and *P2*, respectively (n=39, 34, and 118) (**Fig. 4B, C and Table S2**). The synergy between the injury-responsive activities of *P2* and regeneration enhancers likely contributes to the extremely high rates of injury-dependent expression observed in injured fins. In both hearts and fins, the efficiency of *E1b* associated with the regeneration enhancers was lower than that of *P0.8* or *P2*. Given that *P0.8* and *P2* originate from a regeneration-specific gene, our studies imply that regeneration enhancers can more effectively direct injury-inducible expression when paired with a minimal promoter derived from a regeneration-specific gene.

### Enhancer-promoter pairs display differential eye injury-induced activation

Our F_0_ transgenic assays of *LEN* and *103runx1EN* recapitulated their injury-responsive activities in hearts and fins (Goldman et al., 2017; Kang et al., 2016), indicating the feasibility of our approach for revealing novel enhancer activity. We next applied our transgenic enhancer assays to a new injury model. As the activities of *LEN* and *103runx1EN* in other injured contexts remain unknown, we tested their injury-dependent activities in eyes. To induce eye injury, we passed the tip of a glass needle fully through the right eye of each larva, damaging multiple tissues, including the sclera, retina, and lens (**Fig. 5A**). We first examined the leakiness of the minimal promoters and then assessed the injury-dependent activity of the enhancers. Our assays performed with *E1b*, *P0.8*, and *P2* alone demonstrated no directed expression in uninjured or injured eyes (n=75, 55, and 57, respectively), verifying that all three promoters can be used for enhancer testing (**Fig. 5B, C and Table S2**). Next, we examined the activities of *LEN* and *103runx1EN* in injured eyes, as *lepb* and *runx1* are both induced upon eye injury (Kramer et al., 2021; Zhao et al., 2014). *LEN* drove injury-induced expression in the eye when paired with each tested promoter (21%, n=43 *for E1b;* 49%, n=41 for *P0.8*; 40%, n=35 for *P2*) (**Fig. 5B, C and Table S2**), indicating that *LEN* functions as a regeneration enhancer in eyes. In contrast, *103runx1EN* drove expression only when paired with *P0.8* and *P2* (0%, n=38 *for E1b;* 9%, n=33 for *P0.8*; 20%, n=51 for *P2)* (**Fig. 5B, C and Table S2**). The reduction or lack of injury-induced expression when regeneration enhancers are paired with *E1b* suggests that specific promoter-enhancer interactions are necessary for achieving full enhancer activity. Overall, these results demonstrate that our F_0_ transgenic assays can be used to identify regeneration enhancer activity in a wide range of regenerative contexts.

**Figure 5.**
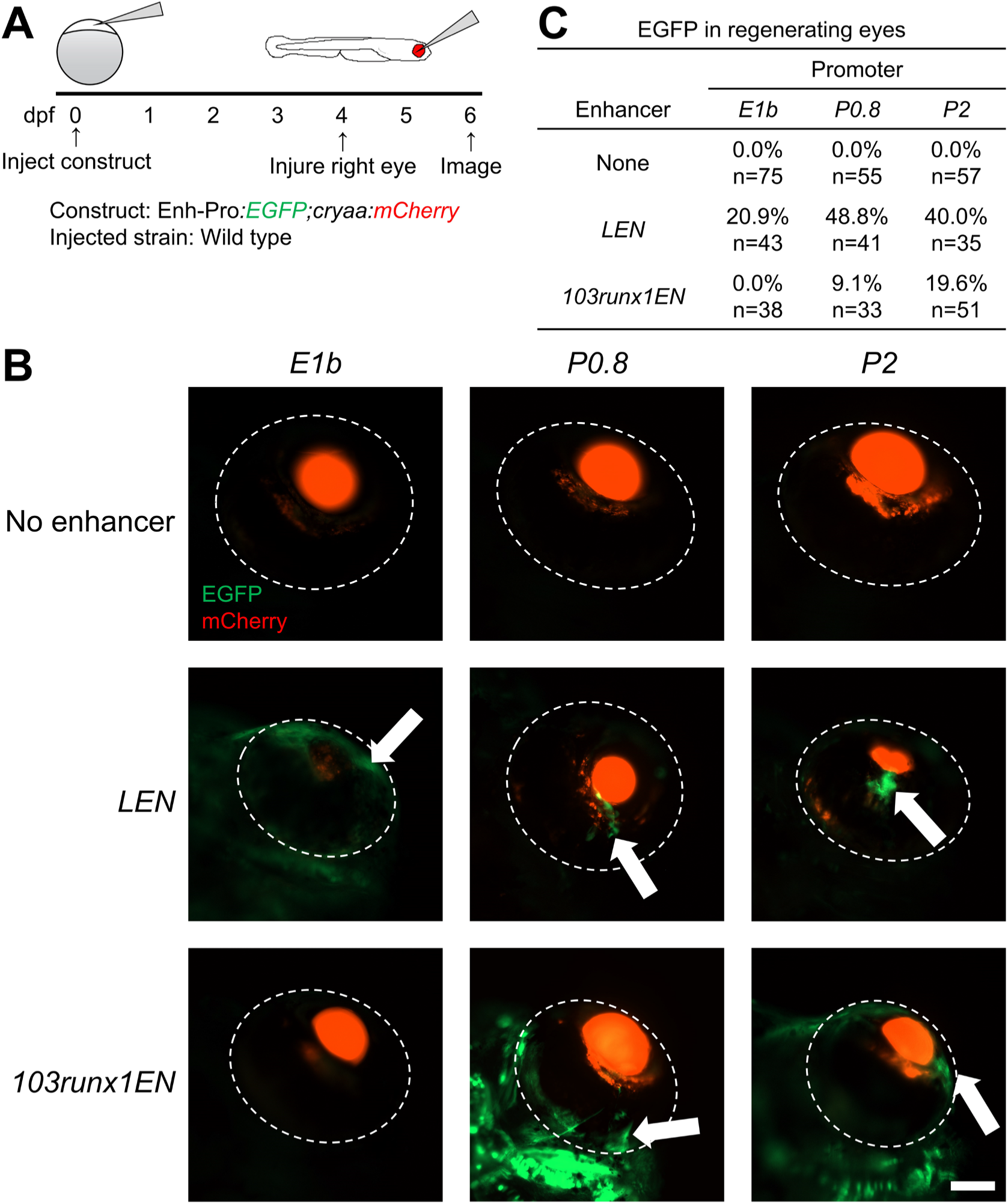
Activity of minimal promoters and regeneration enhancers in injured eyes. (**A**) Experimental design. Transgenic constructs were injected into one-cell-stage wild-type embryos, eyes were injured at 4 dpf, and EGFP expression was observed at 6 dpf. (**B**) 6-dpf F0 regenerating eyes following injury. Arrows indicate EGFP expression. (**C**) Table of larvae displaying eye EGFP expression for each promoter alone or when paired with *LEN* or *103runx1EN*. Scale bar: 100 μm.

### Downstream elements influence enhancer-promoter compatibility

Our assays revealed differences in the efficiency of *E1b* and *P0.8/P2* depending on the enhancer type. *E1b* favors developmental enhancer activity, while *P0.8* and *P2* favor regeneration/injury enhancer activity. To identify potential factors contributing to the higher efficiency of *E1b* when paired with developmental enhancers, we analyzed the sequences within the *E1b* promoter construct. We found that the *E1b* promoter construct used in this and previous studies (Bar Yaacov et al., 2019; Birnbaum et al., 2012; Booker et al., 2013; Hirsch et al., 2018; Kim et al., 2014; Laarman et al., 2019; Li et al., 2010; Oksenberg et al., 2014; Yanovsky-Dagan et al., 2015) is composed of the 42-bp *E1b* minimal promoter and two downstream elements—the 5′ untranslated region (UTR) of the carp *beta-actin* gene and an intron from the rabbit *beta-globin* gene (**Fig. S1A**). These downstream elements provide a transcriptional start site and splicing sites and have been reported to improve transgene activity (Chatterjee et al., 2010; Distel et al., 2009; Horstick et al., 2015; Scheer and Campos-Ortega, 1999), with promoter-proximal splicing sites being known to influence gene transcription (Choi et al., 1991; Furger et al., 2002). Thus, we hypothesized that adding a 5′ UTR and an intron to *P0.8* would improve its efficiency when paired with developmental enhancers without reducing the efficiency of regeneration enhancers. To this end, we generated a zebrafish synthetic promoter (*ZSP*) in which the carp *beta-actin* 5′ UTR and the rabbit *beta-globin* intron were added immediately downstream of *P0.8* (**Fig 6A**). We first assessed the leakiness *ZSP* activity by examining *ZSP* either alone or paired with *LEN* in uninjured developing larvae. *ZSP* displayed low leakiness, driving expression in the uninjured heart, eye, and fin fold in no more than 1.5% of larvae (n=206 and 134 for *ZSP* and *LEN-ZSP*, respectively) (**Fig. 6B, C and Table S2**). These results suggest that *ZSP* activity is tightly regulated in uninjured contexts, indicating its usefulness for enhancer assays.

**Figure 6.**
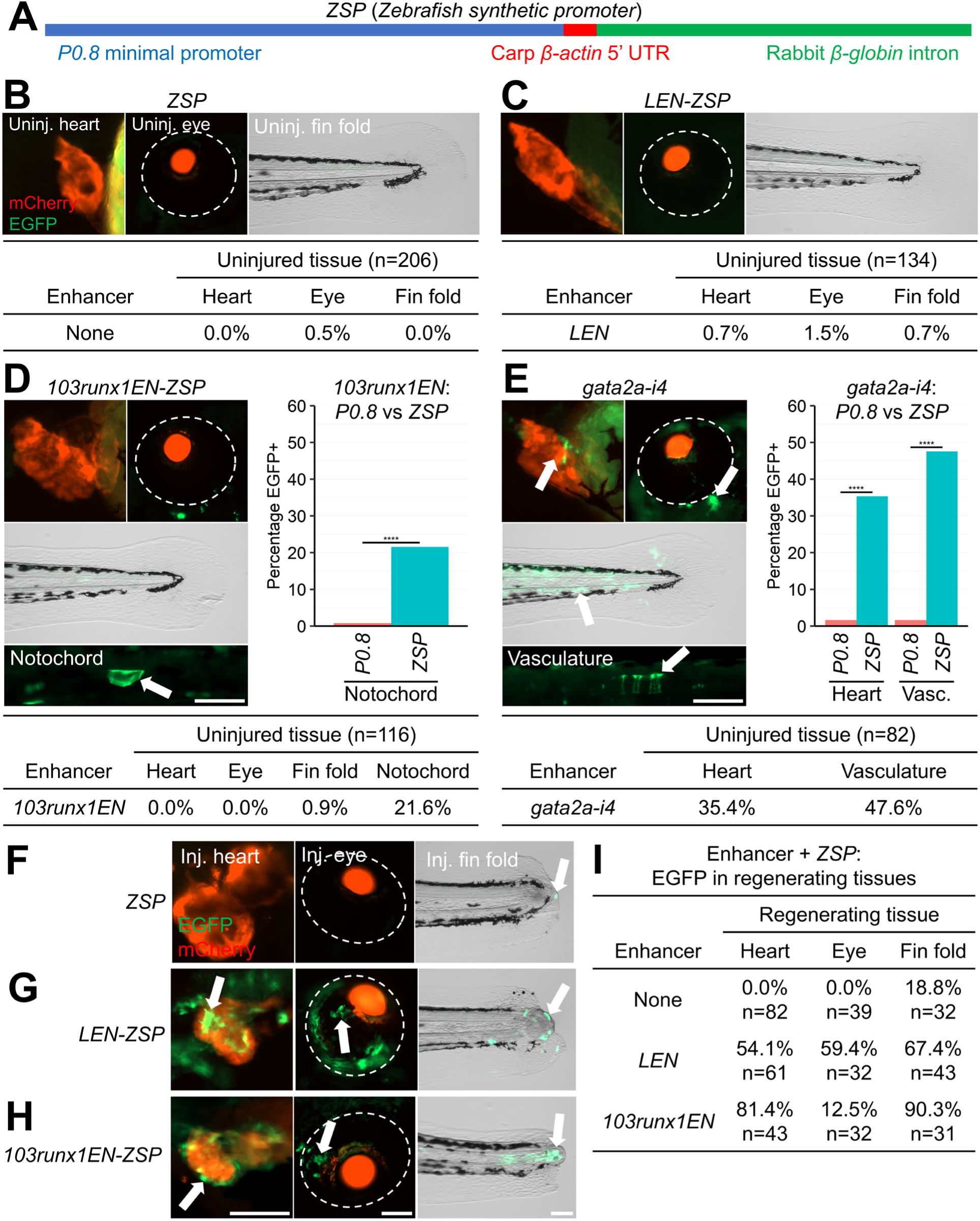
Adding downstream elements to the *P0.8* minimal promoter increases developmental enhancer activity. (**A**) Structure of the *ZSP* synthetic minimal promoter. (**B-E**) 3-dpf F0 uninjured heart, eye, fin fold, notochord, and vasculature of *ZSP* (**B**), *LEN-ZSP* (**C**), *103runx1EN-ZSP* (**D**), and *gata2a-i4-ZSP* (**E**) larvae. Arrows indicate EGFP expression. Eyes are demarcated by dotted lines. The percentage of EGFP^+^ larvae were compared between larvae harboring each *ZSP* construct and each corresponding *P0.8* construct. (**F-H**) 6-dpf F0 regenerating hearts, eyes, and fin folds following injury in *ZSP* (**F**), *LEN-ZSP* (**G**), *103runx1EN-ZSP* (**H**) larvae. (**I**) Table of larvae displaying EGFP expression in each tissue for *ZSP* alone or when paired with *LEN* or *103runx1EN*. Scale bars: 100 μm. ****, p < 0.0001.

We next measured the efficiency of *ZSP* when paired with the developmental enhancers *103runx1EN* and *gata2a-i4*. EGFP was expressed in the notochord in 22% of *103runx1EN-ZSP* larvae (n=116), a significantly higher rate than that of *103runx1EN-P0.8* (**Fig. 6D and Table S2**). Nonspecific expression of *103runx1EN-ZSP* was undetectable in the uninjured heart and eye and occurred in less than 1% of fin folds. EGFP expression was also significantly higher in the hearts and vasculature of *gata2a-i4-ZSP* larvae (35% and 48%, respectively; n=82) than in *gata2a-i4-P0.8* larvae (**Fig. 6E and Table S2**). These data suggest that adding the 5′ UTR and intron sequences downstream of *P0.8* significantly improves its compatibility with developmental enhancer activity.

To determine the efficiency of *ZSP* when paired with regeneration enhancers and its intrinsic activity in injured contexts, we examined the expression of *ZSP*, *LEN-ZSP* and *103runx1EN-ZSP* in hearts, eyes, and fin folds upon injury. While *ZSP* alone directed expression in 19% of injured fin folds (half the rate recorded for *P2*; n=32), it did not drive EGFP in injured hearts and eyes (n=82 and 39, respectively) (**Fig. 6F, I and Table S2**). *LEN-ZSP* drove injury-induced expression in 54% of hearts, 59% of eyes, and 67% of fin folds (n=61, 32, and 43, respectively), showing a significant increase in fin fold expression relative to *LEN-P0.8* (**Fig. 6G, I and Table S2**). The *103runx1EN-ZSP* construct also drove significantly higher rates of EGFP induction in injured hearts and fin folds than *103runx1EN-P0.8* (81%, n=43 for hearts; 90%, n=31 for fin folds), and its activity in injured eyes was similar to that of *103runx1EN-P0.8* (13%, n=32) (**Fig. H, I, Table S2**). These data suggest that the inclusion of downstream promoter elements can improve the efficiency of minimal promoters when paired with developmental enhancers without decreasing their compatibility with regeneration enhancer activity.

Overall, our study demonstrates that both enhancers and promoters play important roles in governing gene expression and that they interact with each other in complex ways. Thus, the characteristics of each enhancer and promoter should be carefully considered when designing constructs for transgenic assays in zebrafish.

## DISCUSSION

Transgenic enhancer assays in zebrafish have been performed using many different minimal promoters (**Table S1**), and it remains unclear how compatibly each promoter performs with various developmental and regeneration enhancers. Here, we assessed the performance of the *E1b*, *P2*, *P0.8*, *ZSP*, *fos*, and DSCP promoters in transgenic enhancer assays in zebrafish. Our study showed that the mouse-derived *fos* and synthetic *Drosophila*-derived DSCP promoters exhibited high levels of leaky expression, displaying ectopic expression in numerous tissues. *E1b*, *P2*, *P0.8*, and *ZSP* drove minimal or no ectopic expression, making them more suitable for use in transgenic screens because enhancer activity is not masked by leaky promoter activity. Developmental enhancer activity appears to exhibit better performance when paired with *E1b*, while injury-responsive enhancer activity works more efficiently with *P0.8* and *P2*, indicating differential enhancer-promoter preference. The synthetic promoter *ZSP*, consisting of *P0.8* together with additional downstream elements, displays a higher efficiency with developmental enhancers than *P0.8* while maintaining its compatibility with injury-responsive enhancer activity. These results imply that elements downstream of the promoter, such as UTRs, splicing sites, and introns, are also important *cis*-regulatory elements affecting transcription.

Based on our findings, we provide guidelines for designing transgenic enhancer assays in zebrafish. First, minimal promoters are crucial regulatory elements that influence gene expression and cannot be considered universal. A previous study in zebrafish investigated combinations of the enhancers and promoters of genes that are active in early development, revealing that the intensity of transgene expression and the level of tissue specificity varied dramatically among different enhancer-promoter combinations (Gehrig et al., 2009). A study in *Drosophila* also reported enhancer-promoter specificity (Zabidi et al., 2015). In the fly, the promoters of housekeeping and developmental genes markedly differ in their characteristics, including their enrichment for specific transcription factor motifs, compositions of core promoter elements, and bound trans-acting factors. A recent high-throughput study in cultured human cells also found distinct categories of promoters (Bergman et al., 2022). Promoters of ubiquitously expressed genes exhibit strong intrinsic activity and low responsiveness to enhancers, while promoters of genes that are variably expressed across cell-types have weak intrinsic activity and high compatibility with enhancers. Because transcriptional specificity is encoded by both enhancers and promoters, appropriate minimal promoters need to be selected when performing transgenic enhancer assays.

Second, a minimal promoter needs to be tailored to the enhancers being investigated. Our study and previous studies have provided evidence that the most appropriate minimal promoter is the cognate promoter from the flanking gene of the enhancer being tested (Quillien et al., 2017). A zebrafish study revealed that endothelial enhancers showed increased expression levels in endothelial cells when paired with their cognate promoters versus a basal promoter (Quillien et al., 2017). In agreement with this finding, *LEN* drove higher rates of injury-responsive expression when paired with its cognate promoter (*P2*, *P0.8*, and the *P0.8*-derived *ZSP*) than with *E1b*. Given these data, in can be inferred that a cognate promoter of an enhancer’s target gene can provide a more robust and accurate readout of the enhancer’s activity in its native context. However, this is unlikely to be feasible in a high-throughput assay. In such a case, we recommend selecting a promoter from a gene that is a well-characterized marker in contexts in which the tested enhancers are predicted to be active. However, examination of the intrinsic activity of the promoter directing ectopic expression is a crucial prerequisite. As shown in a high-throughput study, most promoters appear to possess intrinsic enhancer activity (Nguyen et al., 2016). In our studies, we found that *P2* and, to a lesser extent, *E1b* and *ZSP* could drive injury-induced expression in the absence of an enhancer during the larval stage. Thus, care should be taken to ensure that the promoter does not intrinsically drive expression in the cell types, tissues, or contexts where the enhancer is expected to drive expression.

In summary, our work demonstrates that enhancer-promoter combinations are subject to complex interactions that can drive dramatically different expression patterns during development or regeneration. The proper selection of minimal promoters is important to ensure the accuracy of transgenic assays, and the characteristics and potential compatibilities of each enhancer and promoter should be carefully considered when designing transgenic constructs. We propose that it may be helpful to test multiple minimal promoters or use each enhancer’s cognate promoter to conclusively determine enhancer activity in F_0_ transgenic assays.

### Limitations of the study

Our transgenic approach utilizes random insertions, so the activity of integrated constructs may be influenced by the surrounding regions, known as the position effect. To avoid this, researchers studying mice and *Drosophila* recently established targeted insertion methods to increase reproducibility and decrease chromosomal effects on transgene expression patterns and intensities (Kvon et al., 2014; Kvon et al., 2020). Developing an efficient method for site-directed transgenesis in zebrafish will improve the effectiveness and reproducibility of transgenic enhancer assays.

## METHODS

### Zebrafish maintenance and procedures

Embryos were collected by mating wild-type or transgenic male and female zebrafish from the Ekkwill (EK) strain. Water temperature of adult fish was maintained at 26 °C. Embryos and larvae were maintained at 28 °C in egg water containing 0.3 g/L sea salt, 0.075 g/L calcium sulfate, 0.0375 g/L sodium bicarbonate, and 0.1% methylene blue. To injure hearts with larvae, *cmlc2:mCherry-NTR^pd71Tg^* (Chen et al., 2013) larvae were incubated in 10 mM metronidazole (MTZ) for 24 hours from 3 to 4 dpf to induce CM ablation. To prevent light inactivation of MTZ, larvae were kept in the dark during incubation. Larvae were transferred to fresh egg water following MTZ treatment for recovery. For tail fin fold injury, 4-dpf larvae were first anesthetized in 0.02% tricaine. The tail was amputated using a scalpel at the ventral gap in melanophores at the distal end of the tail. Larvae were transferred to egg water following amputation for recovery. To injure eyes, 4-dpf larvae were anesthetized in 0.02% tricaine and placed on a 1% agarose pad with the right eye facing upwards. The right eye of each larvae was fully penetrated by a glass capillary needle. The left eye was left uninjured as a control. Following injury, larvae were returned to egg water for recovery. Work with zebrafish was performed in accordance with University of Wisconsin–Madison guidelines.

### Plasmid generation

Plasmids were generated through Gibson assembly or restriction enzyme subcloning. Enhancer, promoter, EGFP, and SV40 polyadenylation signal sequences were assembled on the forward strand of each plasmid. To enable identification of larvae carrying transgenic constructs, a reporter cassette containing mCherry driven by the eye lens-specific *alpha crystallin* (*cryaa*) promoter was placed downstream in the reverse complement direction. The construct is flanked by I-SceI meganuclease and *Tol2* transposon sites, enabling transgenesis through either method (Kawakami and Shima, 1999; Thermes et al., 2002). *E1b*, *P0.8*, *P2*, *LEN*, and EGFP sequences were subcloned from previously generated plasmids (Kang et al., 2016). DSCP was subcloned from the pBPGUw plasmid (Addgene plasmid #17575; http://n2t.net/addgene:17575; RRID:Addgene_17575)(Pfeiffer et al., 2008). *103runx1EN* and *gata2a-i4* enhancer sequences were amplified from EK zebrafish genomic DNA. Primers used for subcloning are listed in **Table S3**.

### Transgenesis and imaging

F_0_ transgenic larvae were generated using Tol2 transgenesis (Kawakami et al., 2004). Single cell-stage wild-type or *cmlc2:mCherry-NTR* embryos were injected with a 1 pL of a mixture of 30 ng/μL plasmid DNA, 40 ng/μL Tol2 mRNA, and 0.1% phenol red. Injections were performed using a WPI PV820 microinjector. Larvae were screened for mCherry-positive lenses at 3 dpf. For imaging, anesthetized larvae placed on a 1% agarose pad. Wholemount fluorescent and brightfield images were captured using a Zeiss Axiozoom V16 stereomicroscope. Statistical significance of transgenic construct expression rates was determined via Pearson’s chi-squared test with post hoc pairwise analysis. Significance levels were set to 0.05. Multiple-comparison correction was performed using the Bonferroni correction.

## ACKNOWLEDGMENTS

We thank UW–Madison SMPH BRMS staffs and Kang lab members for zebrafish care. We also thank Dr. Brian Black for constructs. pBPGUw was a gift from Gerald Rubin.

## AUTHOR CONTRIBUTIONS

Biological Experiments: I.J.B., B.E., A.K. Conceptualization: I.J.B, J.K. Writing, Reviewing, Editing: I.J.B., J.K. Funding: J.K.

---

**Figure S1.**
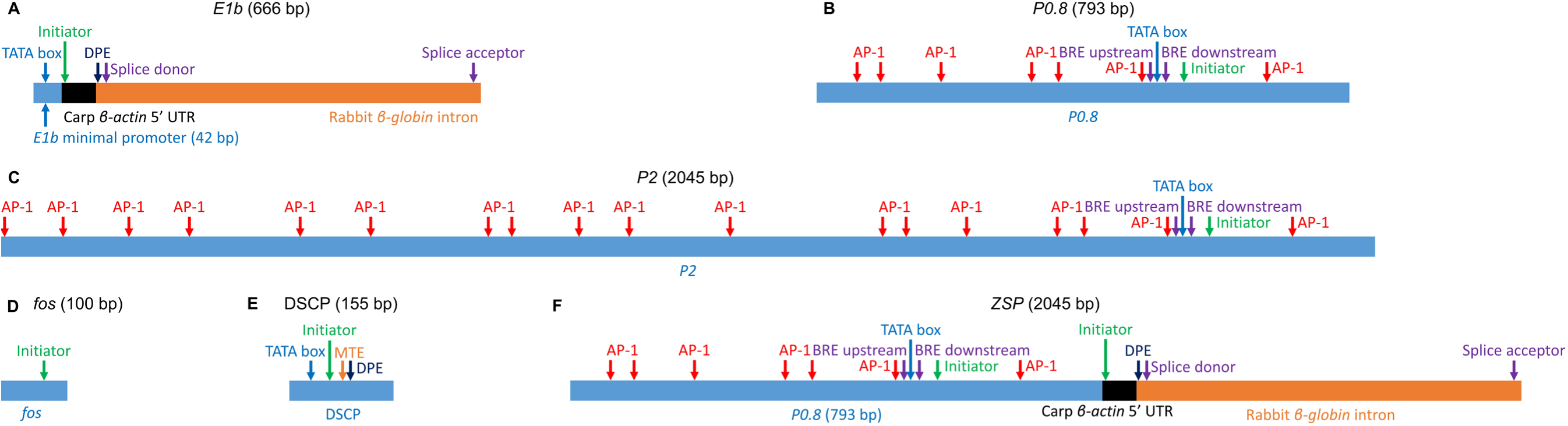
Promoter features of minimal promoter constructs. Promoter features and AP-1 motifs found in the (A) *E1b*, (B) *P0.8*, (C) *P2*, (D) *fos*, (E) DSCP, and (F) *ZSP*. BRE, B recognition element; DPE, downstream promoter element, MTE, motif ten element.

**Figure S2.**
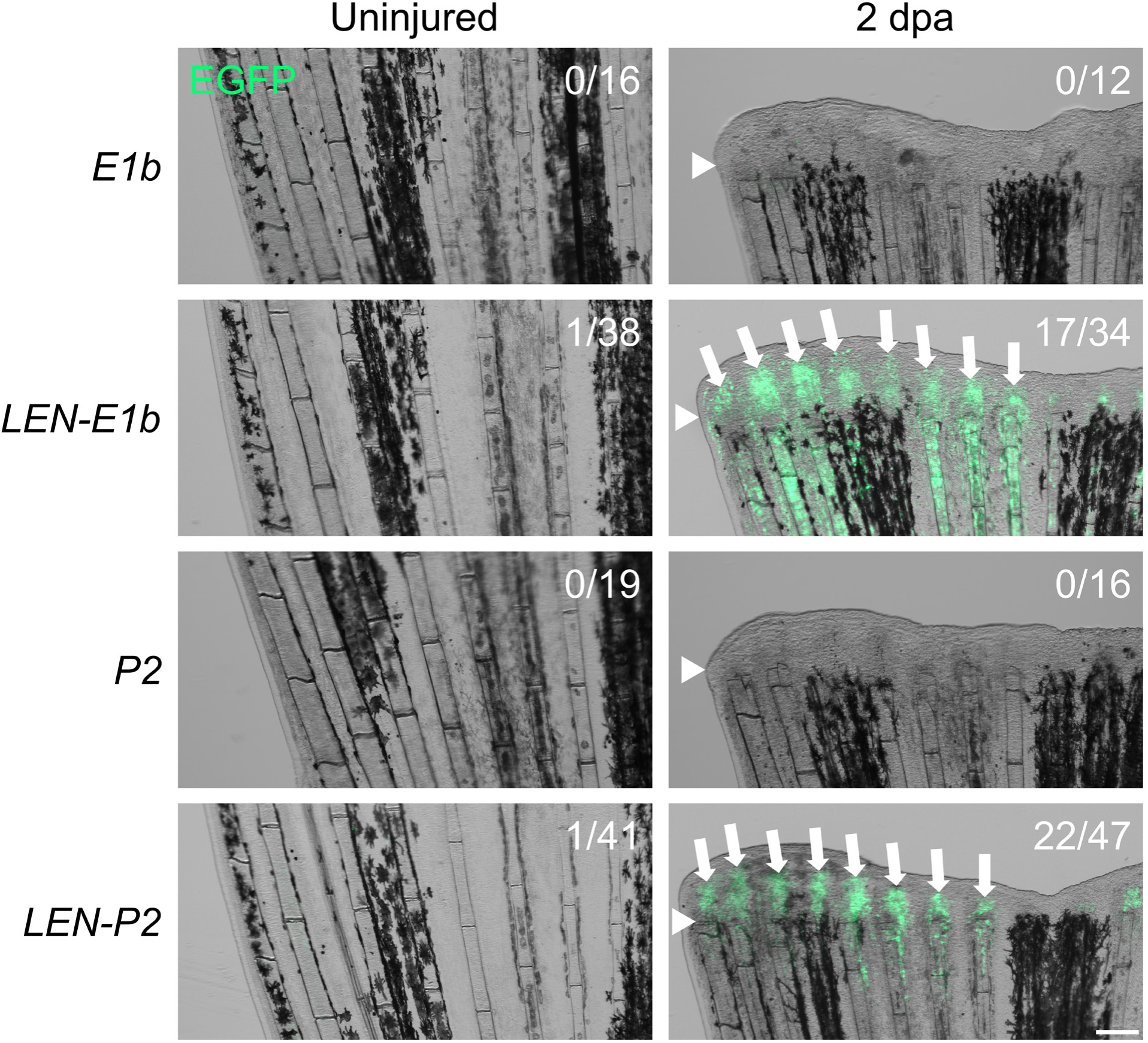
*E1b* and *P2* do not drive injury-responsive expression in adult fins. Uninjured and 2 days post-amputation (dpa) adult F0 *E1b:EGFP*, *LEN-E1b:EGFP*, *P2:EGFP*, and *LEN-P2:EGFP* caudal fins. White arrowheads indicate amputation plane. White arrows indicate injury-induced EGFP expression. Numbers indicate number of fins expressing EGFP. Scale bar: 100 μm.

**Table S1.**
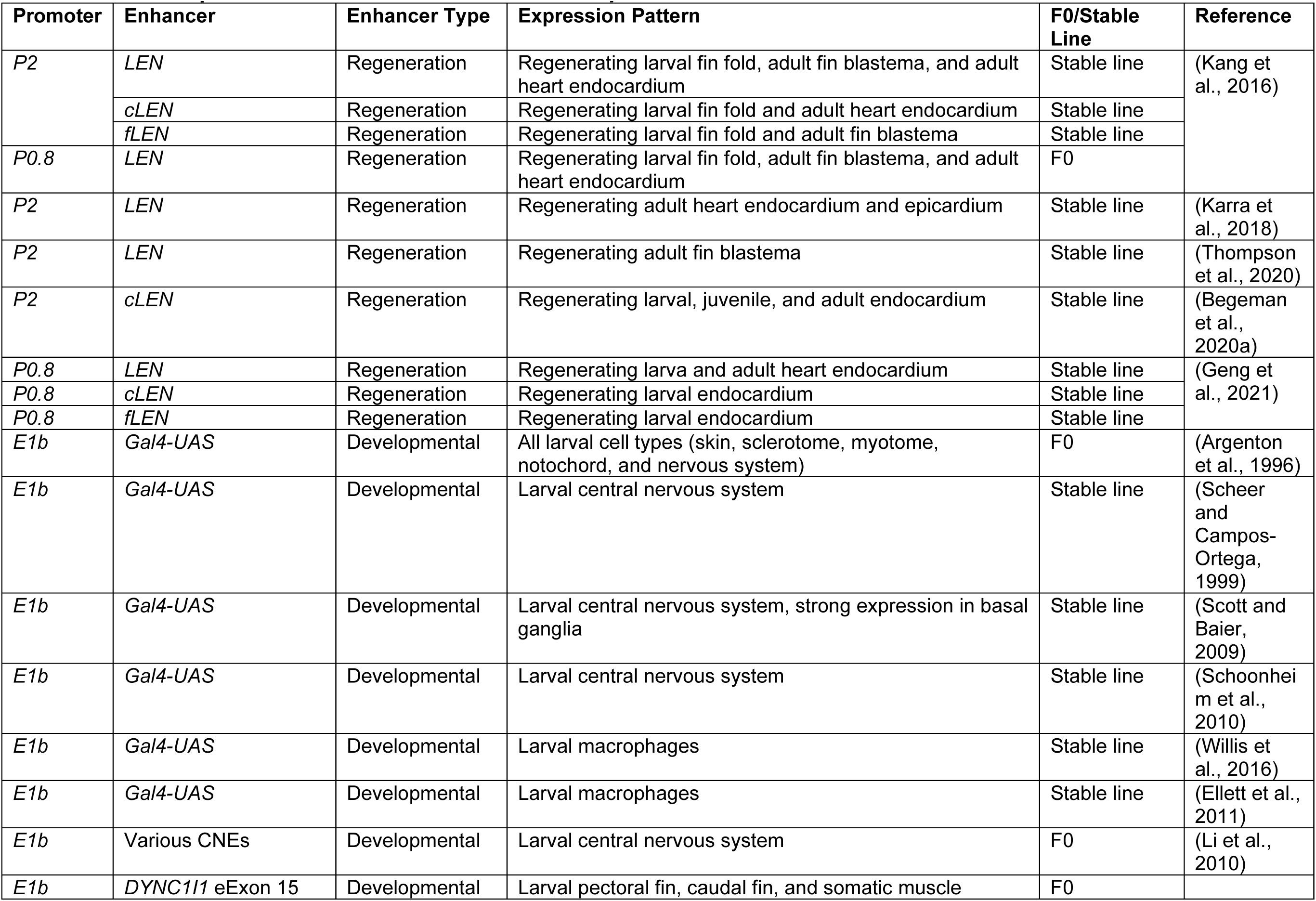
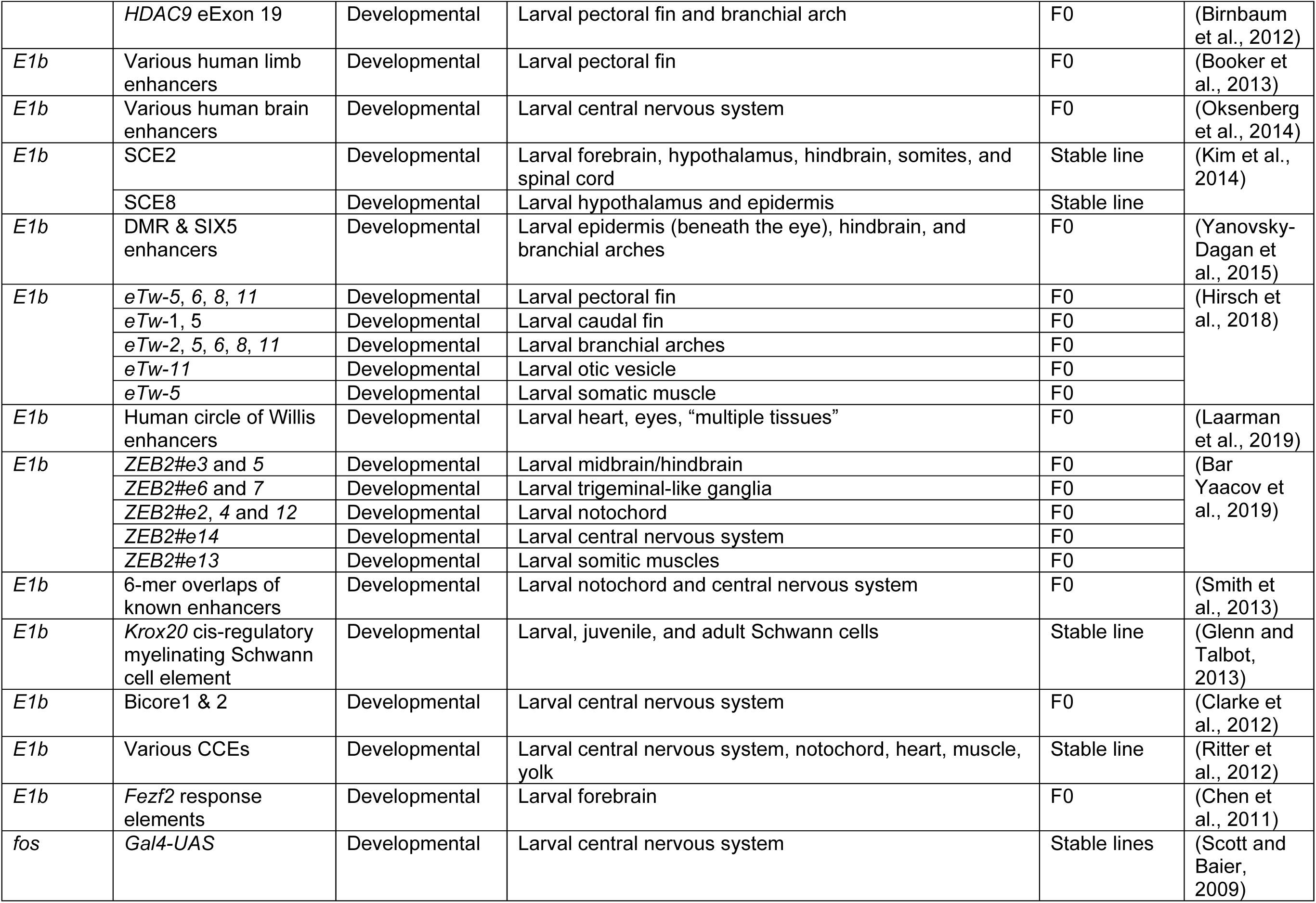
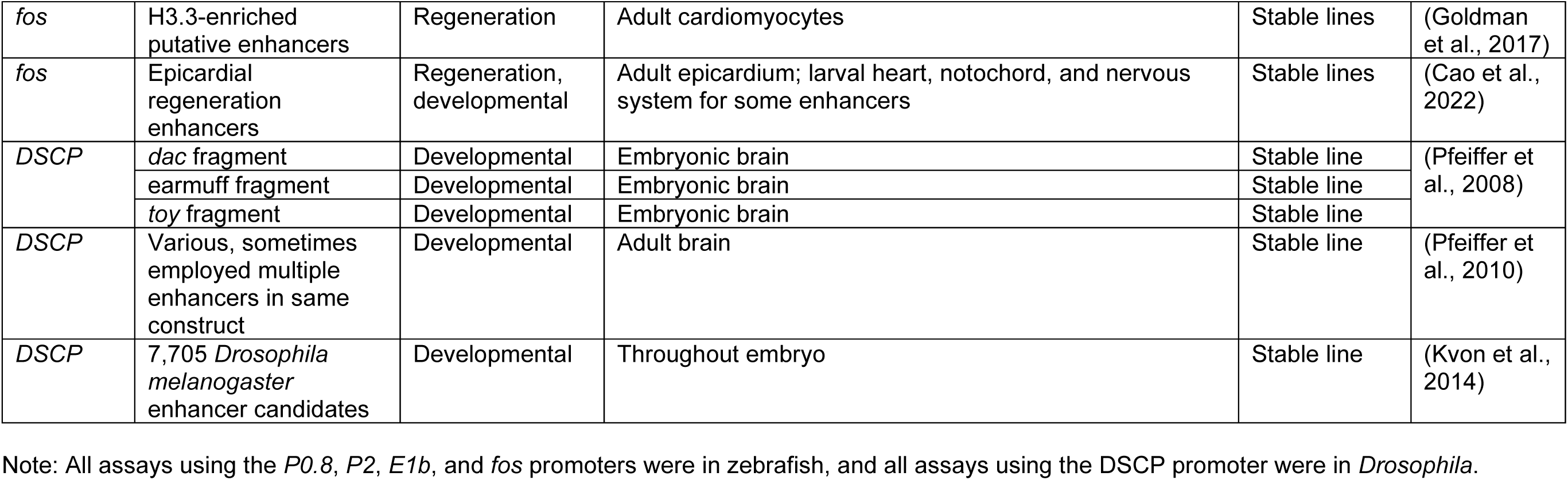
Minimal promoters used in zebrafish and *Drosophila*.

| Promoter | Enhancer | Enhancer Type | Expression Pattern | F0/Stable Line | Reference |
| --- | --- | --- | --- | --- | --- |
| *P2* | *LEN* | Regeneration | Regenerating larval fin fold, adult fin blastema, and adult heart endocardium | Stable line | (Kang et al., 2016) |
|  | *cLEN* | Regeneration | Regenerating larval fin fold and adult heart endocardium | Stable line |  |
|  | *fLEN* | Regeneration | Regenerating larval fin fold and adult fin blastema | Stable line |  |
| *P0.8* | *LEN* | Regeneration | Regenerating larval fin fold, adult fin blastema, and adult heart endocardium | F0 |  |
| *P2* | *LEN* | Regeneration | Regenerating adult heart endocardium and epicardium | Stable line | (Karra et al., 2018) |
| *P2* | *LEN* | Regeneration | Regenerating adult fin blastema | Stable line | (Thompson et al., 2020) |
| *P2* | *cLEN* | Regeneration | Regenerating larval, juvenile, and adult endocardium | Stable line | (Begeman et al., 2020a) |
| *P0.8* | *LEN* | Regeneration | Regenerating larva and adult heart endocardium | Stable line | (Geng et al., 2021) |
| *P0.8* | *cLEN* | Regeneration | Regenerating larval endocardium | Stable line |  |
| *P0.8* | *fLEN* | Regeneration | Regenerating larval endocardium | Stable line |  |
| *E1b* | *Gal4-UAS* | Developmental | All larval cell types (skin, sclerotome, myotome, notochord, and nervous system) | F0 | (Argenton et al., 1996) |
| *E1b* | *Gal4-UAS* | Developmental | Larval central nervous system | Stable line | (Scheer and Campos-Ortega, 1999) |
| *E1b* | *Gal4-UAS* | Developmental | Larval central nervous system, strong expression in basal ganglia | Stable line | (Scott and Baier, 2009) |
| *E1b* | *Gal4-UAS* | Developmental | Larval central nervous system | Stable line | (Schoonheim et al., 2010) |
| *E1b* | *Gal4-UAS* | Developmental | Larval macrophages | Stable line | (Willis et al., 2016) |
| *E1b* | *Gal4-UAS* | Developmental | Larval macrophages | Stable line | (Ellett et al., 2011) |
| *E1b* | Various CNEs | Developmental | Larval central nervous system | F0 | (Li et al., 2010) |
| *E1b* | *DYNC1I1* eExon 15 | Developmental | Larval pectoral fin, caudal fin, and somatic muscle | F0 |  |
|  | *HDAC9* eExon 19 | Developmental | Larval pectoral fin and branchial arch | F0 | (Birnbaum et al., 2012) |
| *E1b* | Various human limb enhancers | Developmental | Larval pectoral fin | F0 | (Booker et al., 2013) |
| *E1b* | Various human brain enhancers | Developmental | Larval central nervous system | F0 | (Oksenberg et al., 2014) |
| *E1b* | SCE2 | Developmental | Larval forebrain, hypothalamus, hindbrain, somites, and spinal cord | Stable line | (Kim et al., 2014) |
|  | SCE8 | Developmental | Larval hypothalamus and epidermis | Stable line |  |
| *E1b* | DMR & SIX5 enhancers | Developmental | Larval epidermis (beneath the eye), hindbrain, and branchial arches | F0 | (Yanovsky-Dagan et al., 2015) |
| *E1b* | *eTw-5*, 6, 8, 11 | Developmental | Larval pectoral fin | F0 | (Hirsch et al., 2018) |
|  | *eTw-1*, 5 | Developmental | Larval caudal fin | F0 |  |
|  | *eTw-2*, 5, 6, 8, 11 | Developmental | Larval branchial arches | F0 |  |
|  | *eTw-11* | Developmental | Larval otic vesicle | F0 |  |
|  | *eTw-5* | Developmental | Larval somatic muscle | F0 |  |
| *E1b* | Human circle of Willis enhancers | Developmental | Larval heart, eyes, "multiple tissues" | F0 | (Laarman et al., 2019) |
| *E1b* | *ZEB2#e3* and 5 | Developmental | Larval midbrain/hindbrain | F0 | (Bar Yaacov et al., 2019) |
|  | *ZEB2#e6* and 7 | Developmental | Larval trigeminal-like ganglia | F0 |  |
|  | *ZEB2#e2*, 4 and 12 | Developmental | Larval notochord | F0 |  |
|  | *ZEB2#e14* | Developmental | Larval central nervous system | F0 |  |
|  | *ZEB2#e13* | Developmental | Larval somitic muscles | F0 |  |
| *E1b* | 6-mer overlaps of known enhancers | Developmental | Larval notochord and central nervous system | F0 | (Smith et al., 2013) |
| *E1b* | *Krox20* cis-regulatory myelinating Schwann cell element | Developmental | Larval, juvenile, and adult Schwann cells | Stable line | (Glenn and Talbot, 2013) |
| *E1b* | Bicore1 & 2 | Developmental | Larval central nervous system | F0 | (Clarke et al., 2012) |
| *E1b* | Various CCEs | Developmental | Larval central nervous system, notochord, heart, muscle, yolk | Stable line | (Ritter et al., 2012) |
| *E1b* | *Fezf2* response elements | Developmental | Larval forebrain | F0 | (Chen et al., 2011) |
| *fos* | *Gal4-UAS* | Developmental | Larval central nervous system | Stable lines | (Scott and Baier, 2009) |
| *fos* | H3.3-enriched putative enhancers | Regeneration | Adult cardiomyocytes | Stable lines | (Goldman et al., 2017) |
| *fos* | Epicardial regeneration enhancers | Regeneration, developmental | Adult epicardium; larval heart, notochord, and nervous system for some enhancers | Stable lines | (Cao et al., 2022) |
| *DSCP* | *dac* fragment | Developmental | Embryonic brain | Stable line | (Pfeiffer et al., 2008) |
|  | earmuff fragment | Developmental | Embryonic brain | Stable line |  |
|  | *toy* fragment | Developmental | Embryonic brain | Stable line |  |
| *DSCP* | Various, sometimes employed multiple enhancers in same construct | Developmental | Adult brain | Stable line | (Pfeiffer et al., 2010) |
| *DSCP* | 7,705 *Drosophila melanogaster* enhancer candidates | Developmental | Throughout embryo | Stable line | (Kvon et al., 2014) |

Note: All assays using the *P0.8*, *P2*, *E1b*, and *fos* promoters were in zebrafish, and all assays using the DSCP promoter were in *Drosophila*.

**Table S2.** Animal number used in this study. Related to Figures 1 and 6.

| Enhancer | Uninjured Tissue |  | Promoter |  |  |  |  |  |
| --- | --- | --- | --- | --- | --- | --- | --- | --- |
|  |  |  | *E1b* | *P0.8* | *P2* | *ZSP* | *fos* | *DSCP* |
| None | Heart | GFP+ | 0 | 0 | 0 | 0 | 14 | 27 |
|  |  | Total | 196 | 57 | 88 | 206 | 134 | 63 |
|  |  | % | 0.0% | 0.0% | 0.0% | 0.0% | 10.4% | 42.9% |
|  | Eye | GFP+ | 0 | 0 | 0 | 1 | 3 | 2 |
|  |  | Total | 196 | 57 | 88 | 206 | 134 | 63 |
|  |  | % | 0.0% | 0.0% | 0.0% | 0.5% | 2.2% | 3.2% |
|  | Fin fold | GFP+ | 0 | 0 | 0 | 0 | 105 | 63 |
|  |  | Total | 196 | 57 | 88 | 206 | 134 | 63 |
|  |  | % | 0.0% | 0.0% | 0.0% | 0.0% | 78.4% | 100.0% |
| LEN | Heart | GFP+ | 0 | 0 | 0 | 1 | 2 | 24 |
|  |  | Total | 227 | 234 | 402 | 134 | 85 | 47 |
|  |  | % | 0.0% | 0.0% | 0.0% | 0.7% | 2.4% | 51.1% |
|  | Eye | GFP+ | 0 | 0 | 0 | 2 | 2 | 0 |
|  |  | Total | 227 | 234 | 402 | 134 | 85 | 47 |
|  |  | % | 0.0% | 0.0% | 0.0% | 1.5% | 2.4% | 0.0% |
|  | Fin fold | GFP+ | 0 | 0 | 0 | 1 | 52 | 45 |
|  |  | Total | 227 | 234 | 402 | 134 | 85 | 47 |
|  |  | % | 0.0% | 0.0% | 0.0% | 0.7% | 61.2% | 95.7% |

Related to Figures 2 and 6
| Enhancer | Uninjured Tissue |  | Promoter |  |  |  |
| --- | --- | --- | --- | --- | --- | --- |
|  |  |  | *E1b* | *P0.8* | *P2* | *ZSP* |
| 103runx1EN | Heart | GFP+ | 0 | 0 | 0 | 0 |
|  |  | Total | 414 | 246 | 166 | 116 |
|  |  | % | 0.0% | 0.0% | 0.0% | 0.0% |
|  | Eye | GFP+ | 0 | 0 | 0 | 0 |
|  |  | Total | 414 | 246 | 166 | 116 |
|  |  | % | 0.0% | 0.0% | 0.0% | 0.0% |
|  | Fin fold | GFP+ | 0 | 0 | 0 | 1 |
|  |  | Total | 414 | 246 | 166 | 116 |
|  |  | % | 0.0% | 0.0% | 0.0% | 0.9% |
|  | Notochord | GFP+ | 61 | 2 | 10 | 25 |
|  |  | Total | 414 | 246 | 166 | 116 |
|  |  | % | 14.7% | 0.8% | 6.0% | 21.6% |
| gata2a-i4 | Heart | GFP+ | 129 | 1 | 0 | 29 |
|  |  | Total | 168 | 61 | 73 | 82 |
|  |  | % | 76.8% | 1.6% | 0.0% | 35.4% |
|  | Vasculature | GFP+ | 159 | 1 | 0 | 39 |
|  |  | Total | 168 | 61 | 73 | 82 |
|  |  | % | 94.6% | 1.6% | 0.0% | 47.6% |

Related to Figures 3 and 6
| Enhancer | Injured Heart | Promoter |  |  |  |
| --- | --- | --- | --- | --- | --- |
|  |  | *E1b* | *P0.8* | *P2* | *ZSP* |
| None | GFP+ | 5 | 0 | 0 | 0 |
|  | Total | 124 | 96 | 109 | 82 |
|  | % | 4.0% | 0.0% | 0.0% | 0.0% |
| *LEN* | GFP+ | 13 | 37 | 58 | 33 |
|  | Total | 74 | 61 | 139 | 61 |
|  | % | 17.6% | 60.7% | 41.7% | 54.1% |
| *103runx1EN* | GFP+ | 13 | 71 | 46 | 35 |
|  | Total | 270 | 148 | 102 | 43 |
|  | % | 4.8% | 48.0% | 45.1% | 81.4% |

Related to Figures 4 and 6
| Enhancer | Injured Fin Fold | Promoter |  |  |  |
| --- | --- | --- | --- | --- | --- |
|  |  | *E1b* | *P0.8* | *P2* | *ZSP* |
| None | GFP+ | 5 | 0 | 21 | 6 |
|  | Total | 124 | 55 | 56 | 32 |
|  | % | 4.0% | 0.0% | 37.5% | 18.8% |
| *LEN* | GFP+ | 11 | 20 | 40 | 29 |
|  | Total | 46 | 45 | 67 | 43 |
|  | % | 23.9% | 44.4% | 59.7% | 67.4% |
| *103runx1EN* | GFP+ | 12 | 24 | 102 | 28 |
|  | Total | 39 | 34 | 118 | 31 |
|  | % | 30.8% | 70.6% | 86.4% | 90.3% |

Related to Figures 5 and 6
| Enhancer | Injured Eye | Promoter |  |  |  |
| --- | --- | --- | --- | --- | --- |
|  |  | *E1b* | *P0.8* | *P2* | *ZSP* |
| None | GFP+ | 0 | 0 | 0 | 0 |
|  | Total | 75 | 55 | 57 | 39 |
|  | % | 0.0% | 0.0% | 0.0% | 0.0% |
| *LEN* | GFP+ | 9 | 20 | 14 | 19 |
|  | Total | 43 | 41 | 35 | 32 |
|  | % | 20.9% | 48.8% | 40.0% | 59.4% |
| *103runx1EN* | GFP+ | 0 | 3 | 10 | 4 |
|  | Total | 38 | 33 | 51 | 32 |
|  | % | 0.0% | 9.1% | 19.6% | 12.5% |

**Table S3.**
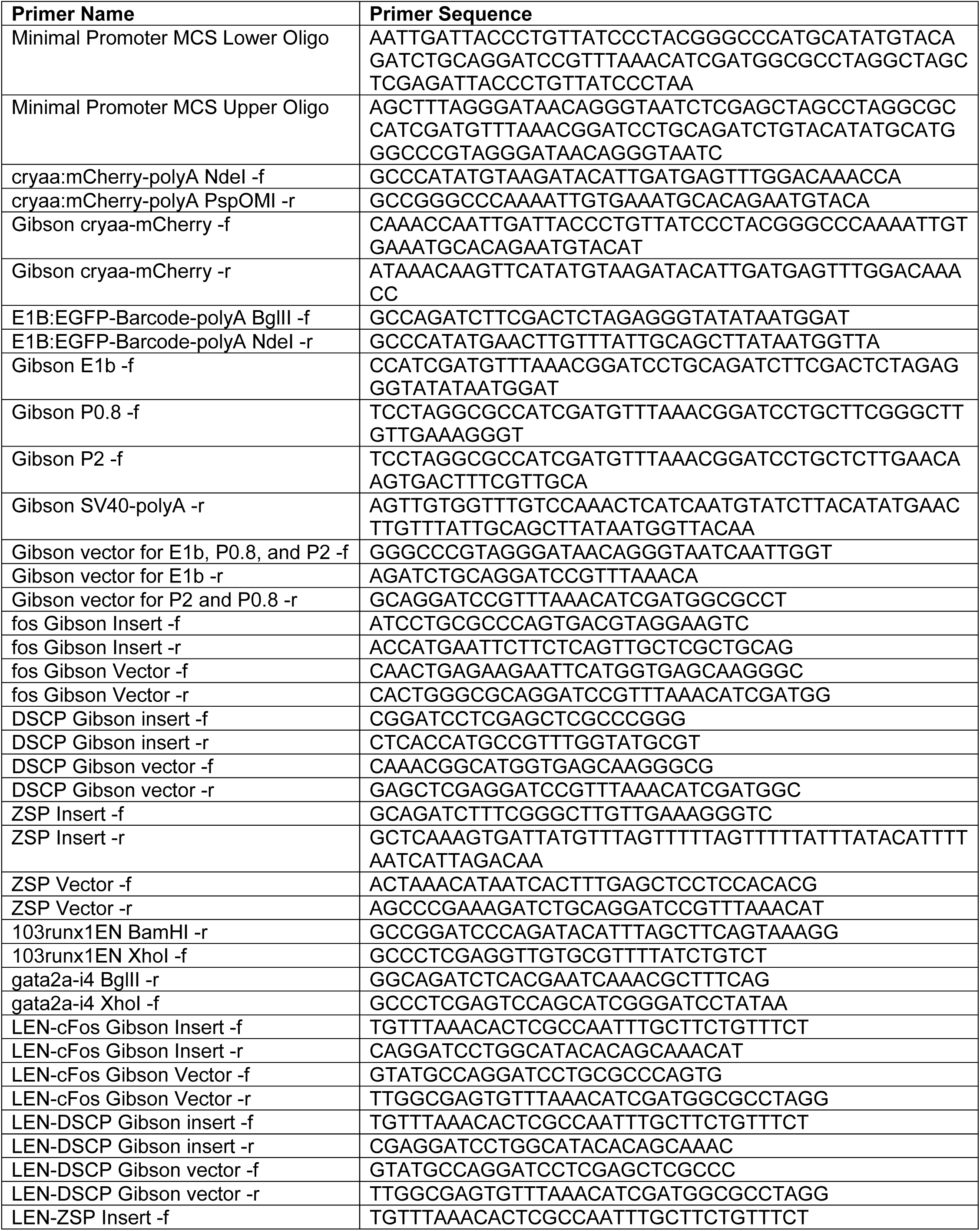
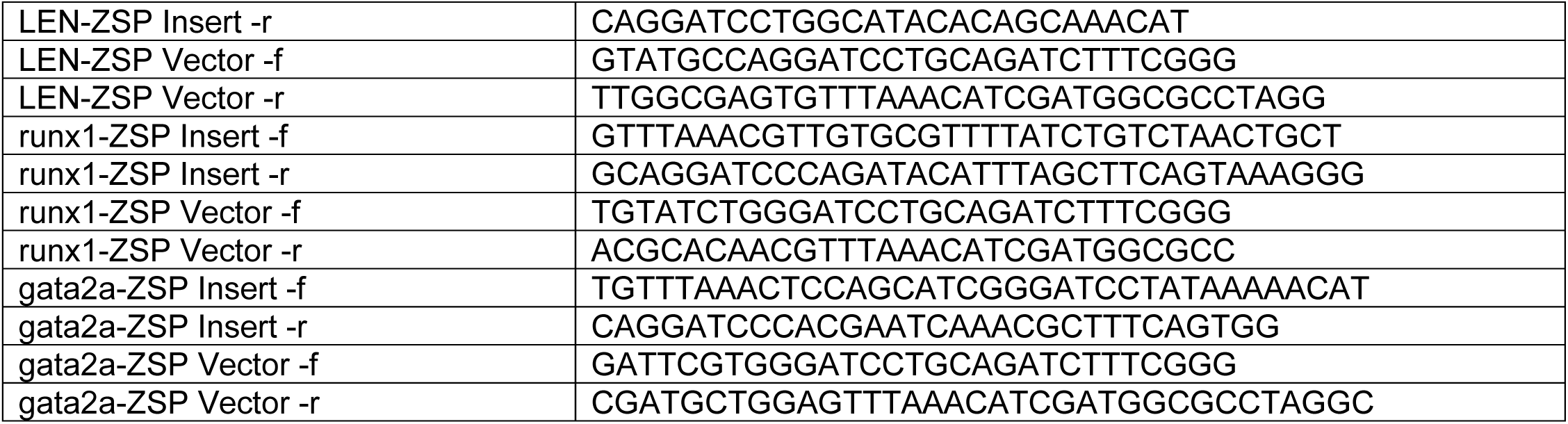
Sequences of primers used in this study.

